# Burning for grassland pollination: recently burned patches promote plant flowering and insect pollinators

**DOI:** 10.1101/2021.08.02.454794

**Authors:** Camila da Silva Goldas, Luciana Regina Podgaiski, Carolina Veronese Corrêa da Silva, Milton de Souza Mendonça

**Affiliations:** Laboratório de Ecologia de Interações, Departamento de Ecologia, Universidade Federal do Rio Grande do Sul, UFRGS, Avenida Bento Gonçalves 9500, Porto Alegre, 91540-000, RS, Brazil

**Keywords:** Pampa, flower visitors, flower resources, fire, resilience.

## Abstract

Grasslands are historically and evolutionarily associated with disturbances, such as fire, that drive biodiversity assembly patterns and biotic interactions. Disturbance suppression in fire-prone ecosystems usually leads to a decline in forb diversity and flowering due to biomass accumulation, which could jeopardize pollinator diversity. In this study, we investigated patterns and drivers of plant flowering and flower insect visitor communities in a chronosequence of patches from different time-since-fire categories in Southern Brazilian grasslands. Old-burnt patches (more than 2 years since fire) had taller vegetation, more biomass and grass cover than intermediate (about 1 year after fire) and freshly-burnt patches (less than 6 months since fire), which had increased bare soil cover. Forb flower abundance was much higher in freshly-burnt patches, directly predicted by the degree of habitat openness. Pollinator insects were then benefited by floral resource aggregation in freshly-burnt patches, increasing in abundance (bees and butterflies) and species richness (bees). Beetle communities were positively influenced by vegetation height. Furthermore, plant species flowering and bee species composition varied between freshly and old-burnt grasslands, with indicator species found for all recovery stages but mainly freshly-burnt patches. Altogether, these results indicate the importance of maintaining freshly-burnt patches in the grassland landscape: it helps to sustain flower diversity, pollination services, and flowering plant reproduction. Our findings support the idea that a mosaic of grasslands from different times-since-fire should be considered for grassland conservation.

## INTRODUCTION

Fire influences global ecosystem patterns and processes, driving vegetation structure and distribution (Bond *et al*. 2005; Bowman *et al*. 2016). Open ecosystems, as grasslands and savannas, evolved in association with fire, hence wildlife and vegetation depend on fire to maintain their diversity and structure (Pausas & Keeley 2009; Keeley *et al*. 2011; Hoetzel *et al*. 2013; Pausas & Parr 2018). Grassland plant species present adaptive traits which create resilience and fitness advantages, such as resprouting after burning (Overbeck & Pfadenhauer 2007; Keeley *et al*. 2011). Moreover, fire removes biomass, opening gaps in vegetation (Fidelis *et al*. 2012), providing space and light access for plants (Lamont & Downes 2011; Fidelis & Blanco 2014). Plant resprouting after fire usually precedes an increase in flowering forbs (Lunt 1994; Aizen & Vazquez 2006; Pyke 2017) benefited by the post-fire environment (e.g., high bare soil, low biomass availability) and low competition with grasses and shrubs (Pickett 1980; Fidelis & Blanco 2014). Forb flowers usually have high-quality resources such as pollen, nectar, oils and resins (Kremen *et al*. 2007) and benefit pollinator communities that depend on them (Potts *et al*. 2003a; Carvalheiro *et al*. 2014). Thus, fire can potentially influence plant attractiveness and pollinator rewards, affecting pollen transfer and plant community reproductive success (Fulton & Carpenter 1979; Potts *et al*. 2001; Van Nuland *et al*. 2013). Furthermore, higher plant community diversity after fire can reflect strongly on animal community dynamics, affecting food chains at all trophic levels and associated ecological processes (Podgaiski *et al*. 2013; Bowman *et al*. 2016).

Pollinators are fundamental to maintaining global vegetation biodiversity (Potts *et al*. 2010). About 88% of angiosperms require pollinating services for reproduction (Ollerton *et al*. 2011). Most animal pollination is carried out thought entomophilia (Ollerton 2017) especially by the four main insect orders Lepidoptera (butterflies and moths), Hymenoptera (bees and wasps), Coleoptera (beetles) and Diptera (flies) (Wardhaugh 2015). Lepidoptera presents most of the flower-visiting species (Ollerton 2017), while bees are considered the most important pollinators on Earth (Potts *et al*. 2010). Nevertheless, habitat loss, fragmentation, and agrochemical use have caused huge pollinator declines, which can negatively impact ecosystems, food production, and human welfare (Mathiasson & Rehan 2020). Actions to maintain pollinator diversity are urgently required (Potts *et al*. 2010). The use of prescribed fires has been indicated as a suitable approach to promote grassland biodiversity, and thus could potentially be applied also to enhance flowering and pollinators (Van Nuland *et al*. 2013; Brown *et al*. 2017; Carbone *et al*. 2019). Nevertheless, fire effects are complex and largely depended on the applied fire regimes and the time after fire considered in monitoring (Keeley *et al*. 2011). For instance, briefly after fire pollinator diversity may increase as a direct response to floral resources, but as time passes diversity usually decreases due to habitat structure and vegetation changes (Potts *et al*. 2003a; Fidelis *et al*. 2014). Understanding how plant flowering and pollinator communities are structured along post-fire recovery in fire-prone ecosystems is an important step to support their conservation and management.

Here we studied flowering plants and insect pollinator communities in a chronosequence of post-fire recovering patches in subtropical natural grasslands. Southern Brazilian grasslands have been subjected evolutionarily and historically to fire disturbances (Pillar & Quadros 1997; Blanco *et al*. 2014). Nevertheless, the role of fire for conservation purposes in these ecosystems has been underestimated (Overbeck *et al*. 2018). A few studies have shown positive patch burning effects on grassland arthropod faunas (Podgaiski *et al*. 2013, 2017; Ferrando *et al*. 2016) and, more recently, on mutualistic networks (da Silva *et al*. 2020; Beal-Neves *et al*. 2020). However, more investigations are necessary to better understand fire-induced changes though time in grassland habitat structure, flowering plants, and specifically on different insect pollinator groups. We first postulate that grasslands from different times-since-fire will differ in habitat structure, i.e., more recently burned patches will present increased bare soil, reduced plant biomass and grass cover in comparison to grasslands remaining longer without fire disturbance (Overbeck *et al*. 2005; Fidelis *et al*. 2012; Podgaiski *et al*. 2014; da Silva *et al*. 2020). Associated with these dynamics, we expect freshly-burnt patches to show high flower abundance and richness of plants flowering due to rapid resprout after fire (Fidelis & Blanco 2014; Pike 2017) which in turn would positively enhance abundance and richness of insect pollinator groups through floral resource improvement (Potts et al. 2003a; Potts et al. 2003b). Furthermore, we hypothesize that species composition of both plant and pollinator species would change along the post-fire chronosequence, with more marked differences between freshly and old-burnt grasslands, due to environmental filtering processes (Vogel *et al*. 2010; de Deus & Oliveira 2016).

## MATERIAL AND METHODS

### Study area

We conducted this study in natural grasslands from the Pampa Biome in the Parque Natural Municipal Saint’ Hilaire (PNMSH) (30°5′ S, 51°5′ W, 82 m a.s.l.), Viamão municipality, Rio Grande do Sul State, Brazil. South Brazilian grasslands (Campos Sulinos - Overbeck *et al*. 2007) are ecosystems very rich in plant species, composed of C4 and C3 grasses, shrubs and a high diversity of forbs and other plant life-forms (Overbeck & Pfadenhauer 2007; Overbeck *et al*. 2007). Representative plant families are Poaceae, Asteraceae, Fabaceae and Rubiaceae (Overbeck *et al*. 2006). These grasslands are disturbance-dependent (fire/grazing; Pillar & Quadros 1997), and disturbance suppression results in changes in vegetation structure and typical biodiversity loss (Oliveira *et al*. 2017; Pillar & Vélez 2010). Grasslands under suppression or low-frequency disturbance regimes usually develop plant communities with high dominance of C4 grasses and shrub encroachment, which increases the risk of catastrophic wildfires due to biomass accumulation (Pillar & Vélez 2010), or forest species encroachment in areas of forest-grassland mosaics (Oliveira & Pillar 2004). Fire has been a frequent disturbance in South Brazilian grasslands since the Holocene (Behling *et al*. 2004; Behling & Pillar 2007). Nowadays, it is traditionally used by farmers, especially in the highland region, as a management tool to remove dry biomass and “renew” pasture after winter (Overbeck *et al*. 2005; Fidelis *et al*. 2010a; Andrade *et al*. 2016).

The PNMSH is located within an urban region, and encompasses about 450 ha of native forest and 300 ha of native grasslands intermingled in a complex mosaic. Cattle grazing does not currently occur, and fire is locally the major grassland disturbance. Grassland burnings are frequent every two or three years from October to February (spring and summer), and are accidentally (e.g., from discarded cigarettes) or intentionally (i.e., arson) induced by the low-income urban population living nearby. Fire intensity is dependent on flammable plant mass accumulation, and it is considered low in these grasslands compared to other fire-prone vegetation types (Fidelis *et al*. 2010b). The burnings, although not controlled, rarely harm the forest ecosystems because of their humid microclimate. The Park rangers from PNMSH record an accurate burning history of each grassland site, and this was used as a basis to select burned patches used in this study. For the study region (including sites within the PNMSH), Beal-Neves *et al*., (2020) have shown a positive relationship between time-since-fire (as an indicative of fire frequency) and urbanization (proximity to urban areas), indicating that grassland patches closer to an urban matrix are more likely to burn. There are no historical records of wildfires, i.e., by natural causes, for the study region.

### Experimental design

We studied 12 burned grassland patches (sampling units). All patches were burned in independent burning events, from three time-since-fire categories: freshly-burnt (FB; n = 4; two to six months after fire), intermediate-burnt (IB; n = 4; about one year after fire), and old-burnt (OB; n = 4; at least two years after fire) (Fig. 1). Half the patches (two of each category) were studied in the 2015/2016 growing season (November to April), and the other half in the 2016/2017 growing season.

**Fig. 1.**
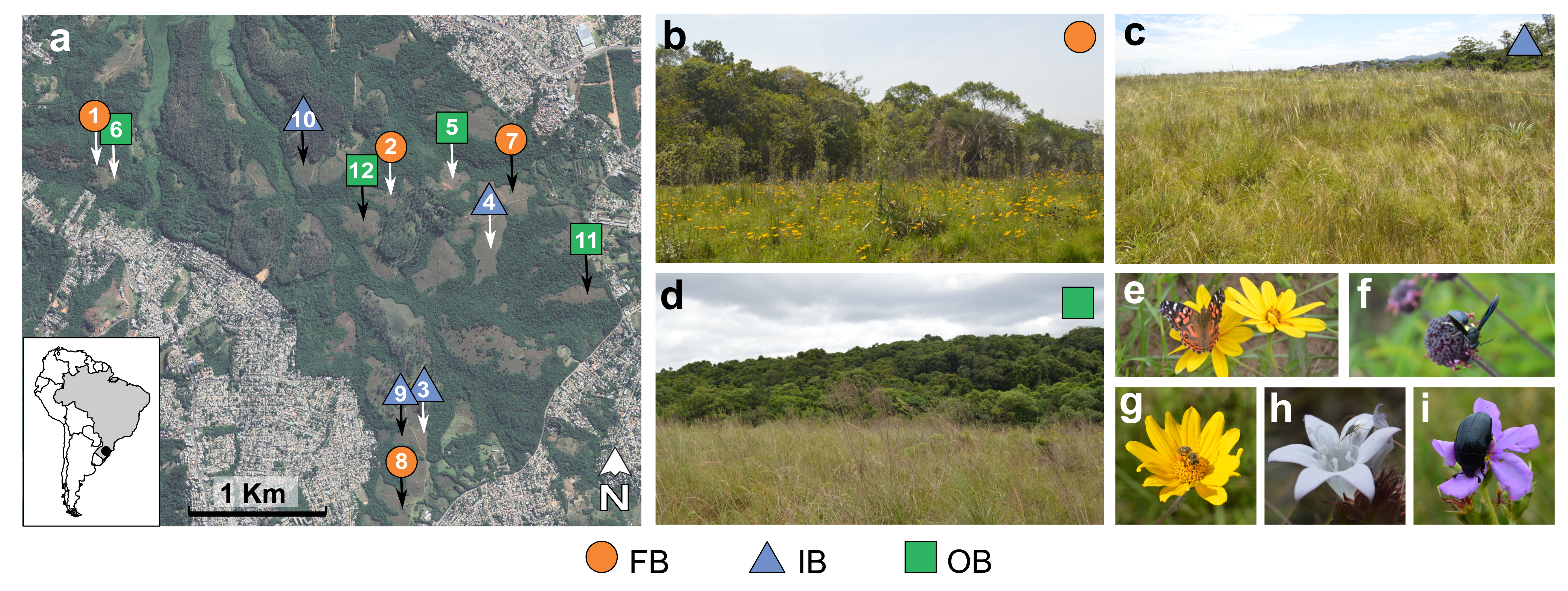
(a) Location of the 12 studied burned patches in the Parque Natural Municipal Saint’ Hilaire in South Brazil. Patches 1 to 6 (white arrows) were sampled in 2015/2016 growing season, and patches 7 to 12 (black arrows) were sampled in 2016/2017. The three time-since-fire categories studied: (b) FB: freshly-burnt (two to six months after fire; orange circle), (c) IB: intermediate-burnt (about one year after fire; light blue triangle), and (d) OB: old-burnt (two years after fire; green square). Insect groups collected in this study: (e) butterfly *Vanessa braziliensis* (Nymphalidae) in *Aspilia montevidensis* (Asteraceae); (f) non-identified wasp (Vespidae) in *Peltodon longipes* (Lamiaceae); (g) bee *Apis mellifera* (Apidae) in *Aspilia montevidensis* (Asteraceae); (h) non-identified fly (Bombyliidae) in *Galianthe fastigiata* (Rubiaceae); (i) beetle *Macraspis dichroa* (Scarabaeidae) in *Tibouchina gracilis* (Melastomataceae).

The selected burned patches were well distributed over the area of the PNMSH (Fig. 1a). The size of the burned patch was estimated based on personal communication with managers of the PNMSH, and further validation with satellite images (when available). It varied from 0.6 (patch 2) to 7.74 ha (patch 4) (average 2.8 ha) (Table S1 - Supporting Information). Proximity to urban areas (i.e., the distance from the center of each patch to the edge of the nearest urban area) varied from 120 m (patch 1) to 1082 m (patch 2) (average 500 m). Grassland proportion in a buffer of 300 m radius around each patch varied from 21.4% (patch 11) to 60.2% (patch 8) (average 39.7%). The complementary proportion represented mostly forests (Table S1 - Supporting Information). Within the same sampling season, all patches were more than 100 m apart from each other, except patches 1 (freshly-burnt) and 6 (old-burnt) that were closer than 20 m (Fig. 1). Despite the high proximity that could facilitate flying arthropod dispersal, habitat preferences of the organisms should prevail because the patches belonged to different time-since-fire categories (Swengel 2001).

**Table 1.** IndVal index and p-values for flowering plant and pollinator species related to time- since-fire categories. FB: freshly-burnt (two to six months); IB: intermediate-burnt (about one year); OB: old-burnt (two years after fire)

| Species | Taxonomic classification | IndVal | p-value | Time-since-fire |
| --- | --- | --- | --- | --- |
| **Plants** |  |  |  |  |
| *Galactia marginalis* (Benth. ex Benth. & Hook. f.) | Fabaceae | 1.00 | 0.007 | FB |
| *Stevia* sp. | Asteraceae | 0.98 | 0.007 | FB |
| *Peltodon longipes* (Kunth. ex Benth.) | Lamiaceae | 0.95 | 0.04 | FB |
| *Lippia* sp. | Verbenaceae | 0.90 | 0.01 | FB |
| *Dyckia leptostachya* (Baker) | Bromeliaceae | 0.84 | 0.05 | IB |
| *Eryngium pristis* (Cham. & Schltdl.) | Apiaceae | 0.93 | 0.03 | FB+IB |
| *Croton gnaphalii* | Euphorbiaceae | 0.98 | 0.01 | OB |
| **Bees** |  |  |  |  |
| *Bombus morio* | Hymenoptera | 1.00 | 0.009 | FB |
| *Augochlorella acarinata* | Hymenoptera | 0.93 | 0.02 | FB+IB |
| **Flies** |  |  |  |  |
| Syrphidae sp. 6 | Diptera | 0.93 | 0.03 | FB+IB |

On each patch, we established six 10 x 10 m plots, distant more than 10 m from each other, where we sampled data related to (i) habitat; (ii) flowering plant communities, and (iii) flower-visiting insect communities.

### Habitat descriptors

We used Hobo® data loggers aboveground (about 15 cm high) to measure air temperature (°C) and relative air moisture (%) of each grassland patch. The sensors recorded climatic variables every 5 minutes during 24 hours at the center of each patch, at the same time in each growing season.

We randomly delimited one quadrat of 1 x 1 m within each plot where we visually estimated cover (%) of bare soil and grasses. We measured vegetation height from ground level to the tallest plant using a tape at five points in each quadrat. After, we mowed the total aerial biomass at ground level inside the quadrat with hedge clippers. The biomass was oven-dried at 60 °C during 72h and weighted in the lab. We sampled these vegetation cover variables twice a year (late November and early April) and in different quadrats (total 12 quadrats per burned patch).

### Flowering plants

Surveys of plants currently blooming and insect pollinators were carried out in three rounds per growing season (i.e., spring: November/December, summer: January/February, and autumn: March/April). We recorded all flowering plant species, and the number of flowers or inflorescences (according to each plant species characteristics) in three randomly assigned parallel parcels of 1 m x 10 m inside each plot. We excluded Poaceae from our observations, since they are generally anemophilous (Faegri and van der Pijl 1979). We collected plant species vouchers, which were identified by local botanists to species or morphospecies (henceforth species).

### Pollinators

We observed each flowering plant species per burned patch during fifteen minutes in three different periods of the day (8-11 a.m., 11 a.m.-2 p.m., and 2-5 p.m.) searching for pollinators. For each plant species, we sampled with entomological nets all foraging flower insect visitors that contacted anthers or flower stigmas (i.e., potential pollinators). Insects were killed in plastic jars with ethyl acetate, and at the lab they were sorted into major groups: bee (Hymenoptera), wasp (Hymenoptera), beetle (Coleoptera), butterfly (Lepidoptera) or fly (Diptera). Individuals were pinned and identified to species or morphospecies (henceforth species). Biological specimens are deposited in the reference collection of Laboratório de Ecologia de Interações, at Universidade Federal do Rio Grande do Sul, UFRGS (Brazil).

### Statistical analyses

All analyses were conducted in R (R Core Team 2020). We first checked for potential relationships between time-since-fire categories and landscape variables from the study site with simple linear effect models. The size of the burned patch (F2,9 = 0.33; p = 0.72), the proximity to urban areas (F2,9 = 0.16; p = 0.85), and the percentage of grassland cover in a buffer of 300 m radius (F2,9 = 2.78; p= 0.11) did not significantly vary with time-since-fire categories (Table S1 - Supporting Information). Additionally, to check for possible structured spatial patterns of pollinator communities in the studied site, we performed Moran’s test in R package *ape* (Paradis & Schliep 2019). We did not detect positive autocorrelation between total pollinator abundance (Moran’s I = -0.46; p= 0.97) or richness (Moran’s I = -0.15; p= 0.63) with the geographic distance of patches. Thus, we conducted data analysis considering no relationships of time-since-fire categories with other landscape variables that could also be affecting pollinator communities, and also with no bias regarding the proximity of the patches in the study site. Generalized linear effect models (GLMs) were further fit always considering the sampling season identity (two levels: 2015/2016 or 2016/2017 growing season) as blocking factor in the models (i.e., first term) to control for possible effects of this variable. Null models included only sampling season as a predictor (i.e., dependent variable ∼ season).

### Habitat structure

We investigated the responses of habitat descriptors to time-since-fire categories with GLMs (i.e., dependent variable ∼ season + time-since-fire category). For each burned patch, temperature and moisture values were averaged from 8 a.m. to 6 p.m., and % bare soil, % grasses, plant biomass, and plant height were averaged per quadrat (m^2^). We fitted Gaussian distribution for models with temperature, moisture, vegetation height and biomass (log transformed) with the R package *stats*, and beta distribution for models considering % grasses and % bare soil with the R package *betareg* (Cribari-Neto & Zeileis 2010). We then performed likelihood-ratio tests comparing these models with null models. Model residuals were evaluated for overdispersion. When time-since-fire was a significant predictor, pairwise differences among the categories were tested using Tukey’s post-hoc test with R package *multcomp* (Hothorn *et al*. 2008). We adjusted p-values for multiple comparisons with false discovery rate correction (FDR; Benjamini & Hochberg 1995).

### Flowering plants

We investigated how the richness of plants flowering, and flower abundance responded to time-since-fire categories with GLMs, as previously described for habitat structure. For each burned patch, the richness of plants flowering was the total number of plant species flowering counted in all parcels within plots (over 180 m^2^) and seasonal rounds (three rounds). Flower abundance was the total number of flowers/inflorescences recorded in all parcels within plots and rounds. We fitted negative binomial distributions and loglink functions for these models with the R package *mass* (Venables & Ripley 2002).

For the response variables that significantly varied with time-since-fire we tested for direct effects of the fire-induced habitat changes with GLM. First, we fit full model with all predictors (additive model: % bare soil, % grasses, plant biomass, and plant height), and then tested for collinearity among predictors using variance inflation factor (VIF) values from the R package *car* (Fox & Weisberg 2019). Two predictors (biomass and % grasses) had VIF values above the recommended upper threshold value of 3 and were excluded from the model (Zuur *et al*. 2010). Using this criterion, we then fitted the model with season + % bare soil + plant height as additive predictor variables (Table S2 – Supporting Information). We calculated pseudo-R-squared values for the models indicating how well the model explains the data in comparison to the null model.

To explore the composition of flowering plant species in the grassland patches from different time-since-fire categories, we generated Principal Coordinate Analysis (PCoA) scatter plots using the function *cmdscale* from the R package *stats*. We used the matrix Wpi, where each patch is described by the incidence of each flowering plant species considering all the plots, and rounds. Singletons were removed. We used the Jaccard index as a measure of community similarity. We additionally added a biplot with habitat structure variables to the ordination, and explored the relationships of each variable with the firsts PCoA axes with Pearson’s correlation. To test whether there were significant differences in species composition between grasslands from different time-since-fire categories we used PERMANOVA with 9999 permutations in the R package *vegan*.

To identify plant species associated to a given time-since-fire category, we calculated the IndVal index, proposed by Dufrêne & Legendre (1997). The IndVal index is maximum (100%) when all individuals from a species, independent of the other species relative abundance, are found in a single category (i.e., high exclusivity) and when the species occurs in all sites for that category (i.e., high fidelity). We used matrix Wpa where each patch is described by the total abundance of each flowering plant species in all plots, and rounds. We used 9999 permutations to test the significance of the IndVal index in the R package *indicspecies* (De Cáceres & Legendre 2009).

### Pollinators

We investigated how pollinator abundance and species richness of each insect group (analyzed separately) responded to time-since-fire categories with GLMs, as previously described for habitat structure and flowering plant variables. For each burned patch, the abundance and richness of pollinator groups represented the total number of individuals or species sampled in all periods of the day, and seasonal rounds. We fitted negative binomial distributions and loglink functions for all models with pollinator response variables.

For those pollinator variables that significantly varied with time-since-fire we searched for direct effects from local habitat structure and flower abundance with GLMs as already described. We checked VIF among the predictor variables (additive model: flower abundance, % bare soil, % grasses, plant biomass, plant height; Table S2 – Supporting Information), and the models were then fitted only with season + flower abundance + vegetation height as predictors.

To evaluate pollinator species composition responses to time-since-fire categories, we performed PERMANOVA (9999 permutations) for each visitor group, as previously described for plant communities. For that, we used matrix Wvi, that describes patches by the incidence of pollinator species, considering all periods of the day and rounds, and the Jaccard index as a measure of community similarity. Singletons were removed. For the insect groups with species composition significantly varying with time-since-fire, we performed PCoAs. We added a biplot with habitat structure and flowering plant variables to the ordination, and explored the relationships of the variables with the PCoA axes and time-since-fire categories. Additionally, we estimated the correlation between plant and pollinator composition matrices with a Mantel test.

Finally, we searched for pollinator indicator species of time-since-fire categories by calculating IndVal index, as already described. For that, we used matrix Wva, that includes the total abundances of each pollinator species, in all periods of the day, and rounds.

## RESULTS

### Habitat structure

Time-since-fire categories differed regarding relative air temperature (χ² = 9.94, df = 2, p < 0.01; Fig. 2a and 2b) and moisture (χ² = 7.79, df = 2, p = 0.02; Fig. 2c and 2d). Freshly and old-burnt grasslands contrasted mostly between 8 a.m to 6 p.m. In this period, freshly-burnt grasslands presented higher air temperature (mean: 34.1°C) and lower moisture (mean: 61.2%) than old-burnt grasslands (mean: 28.1°C; 76.3%).

**Fig. 2.**
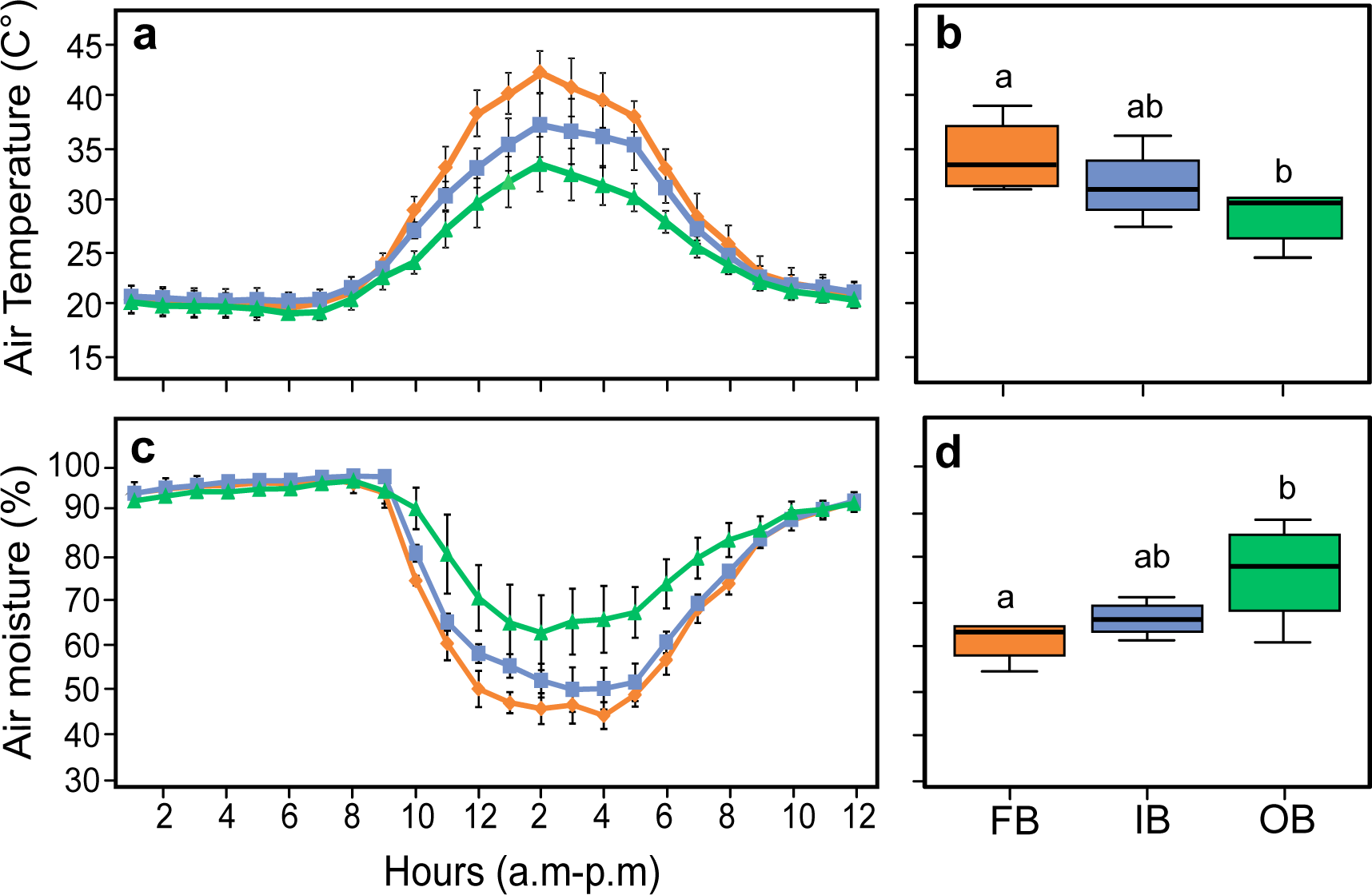
(a) Air temperature (C°) variation during 24 hours, and (b) boxplot of air temperature values between 8 a.m and 6 p.m in grasslands from different time-since-fire categories. (c) Relative air moisture (%) variation during 24 hours, and (d) boxplot of air moisture between 8 a.m and 6 p.m in grasslands from different time-since-fire categories. FB: freshly-burnt (two to six months after fire; orange circle); IB: intermediate-burnt (about one year after fire; light blue triangle); OB: old-burnt grasslands (two years after fire; green square). In b and d, different letters denote significant differences between categories (p ≤ 0.05) tested with GLMs and Tukey test a posteriori. Boxplots represent the median (solid black line), first and third quartile (box limits), and lines denote maximum and minimum values.

Vegetation height was shorter in freshly-burnt (mean: 17.6 cm) and intermediate-burnt (mean: 19.8 cm) than in old-burnt grasslands (mean: 38.4 cm) (χ² = 12.73, df = 2, p = 0.001; Fig. 3a). Aboveground plant biomass was higher in old-burnt (mean: 222.8 g/m^2^) than in intermediate (mean: 98.7 g/m^2^) and freshly-burnt (mean: 124.1 g/m^2^) (χ² = 11.85, df = 2, p = 0.002; Fig. 3b). Grass cover was also higher in old-burnt (mean: 85.2%) than in intermediate (mean: 60.5%) and freshly-burnt (mean: 44.7%; χ² = 16.06, df = 2, p < 0.001; Fig. 3c). Meanwhile, bare soil proportion was higher in freshly-burnt (mean: 22.5%) than in intermediate (mean: 3%) and old-burnt grasslands (mean: 1%) (χ² = 28.93, df = 2, p < 0.001; Fig. 3d).

**Fig. 3.**
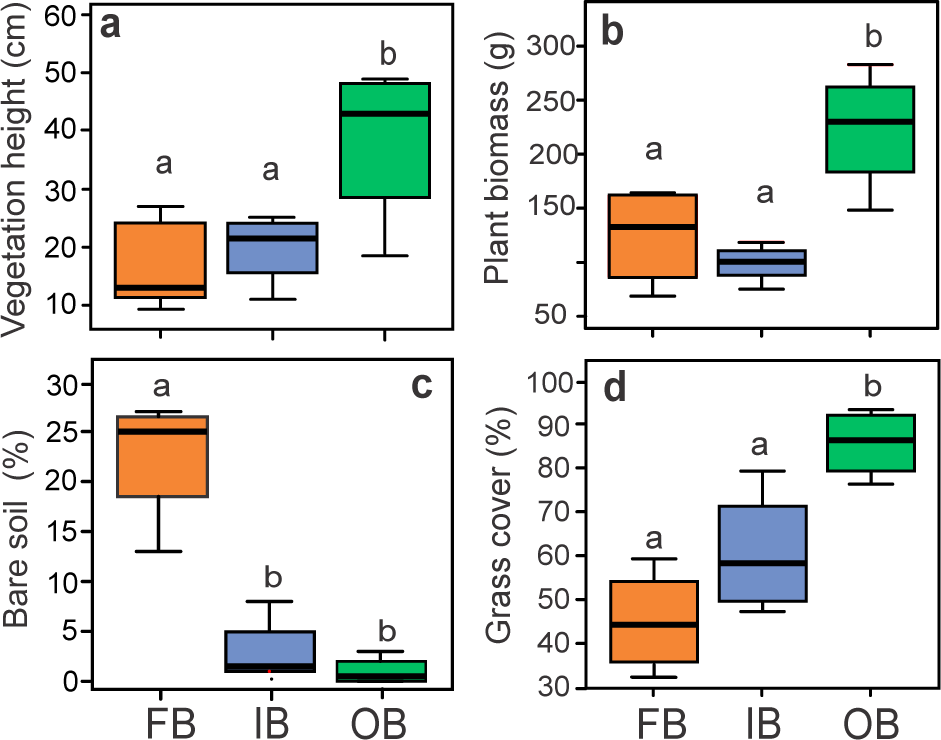
Boxplot of habitat descriptors values: (a) vegetation height, cm; (b) biomass, g; (c) bare soil, %; (d) grass cover, % in grasslands from different time-since-fire categories. FB: freshly- burnt (two to six months after fire; orange); IB: intermediate-burnt (about one year after fire; light blue); OB: old-burnt (two years after fire; green). Different letters denote significant differences between categories (p ≤ 0.05) tested with GLMs and Tukey test a posteriori. Boxplots represent the median (solid black line), first and third quartile (box limits), and lines denote maximum and minimum values.

### Flowering plants

We counted 107 species belonging to 26 plant families (only plants producing flowers; Table S3 - Supporting Information). The most abundant families were Asteraceae with 24% of the total community, followed by Rubiaceae (22%) and Fabaceae (21%). The richest families were Asteraceae (36 species), Fabaceae (16 species), and Apiaceae and Iridaceae with 6 species each.

Species richness of plants producing flowers did not significantly vary among time-since-fire categories (χ² = 3.59, df = 2, p = 0.16), despite higher mean richness values in freshly (mean: 43.2 species) than in intermediate (mean: 35.7 species) and old-burnt (mean: 36.2 species). Flower abundance indeed differed (χ² = 19.96, df = 2, p < 0.001), and was significantly higher in freshly-burnt (mean: 12,155 flowers) than in intermediate (mean: 5,465) and old-burnt grasslands (mean: 2,819 flowers) (Fig. 4a). Both bare soil percentage (deviance = 40.31, df = 9, p < 0.001, pseudo-R^2^ = 66.6%) and vegetation height (deviance = 9.42, df = 8, p < 0.002, pseudo-R^2^ = 62.1%) explained flower abundance. As bare soil percentage increased, abundance of flowers also increased (Fig. 4b). On the contrary, as vegetation height increased, abundance of flowers decreased (Fig. 4c).

**Fig. 4.**
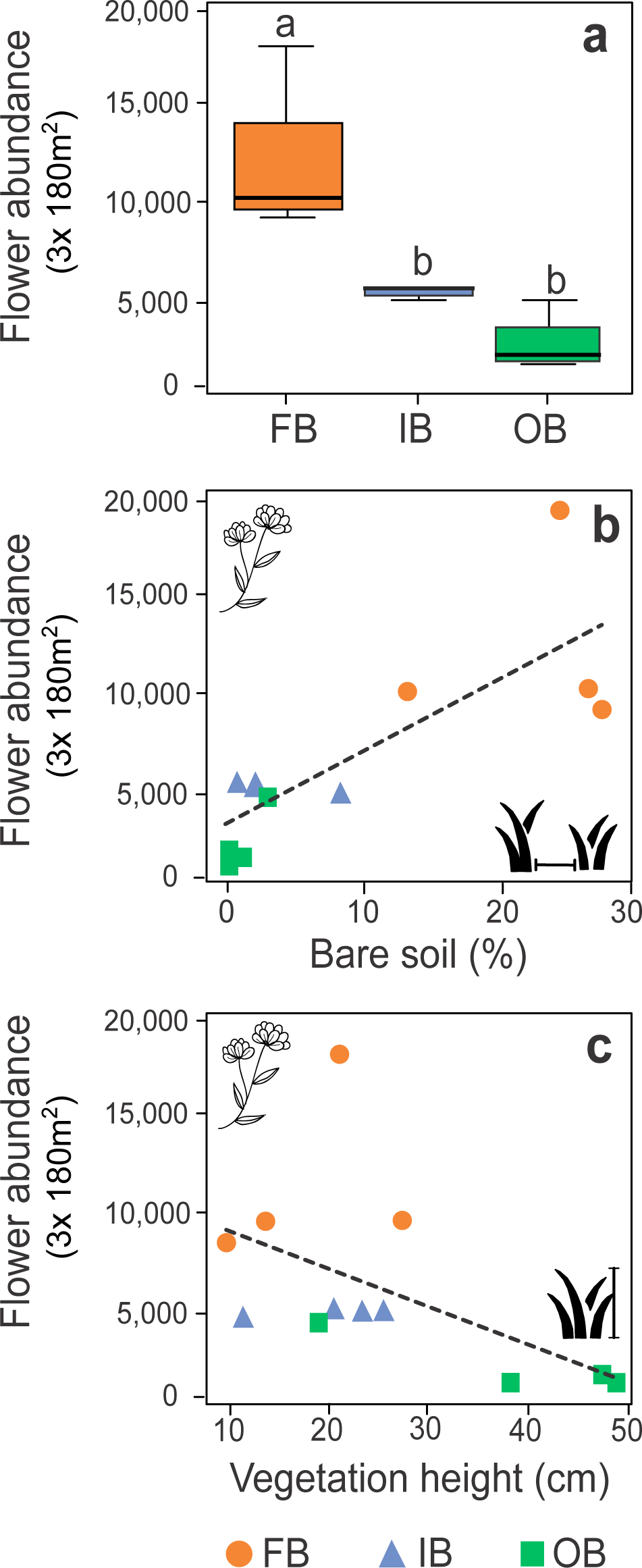
(a) Boxplot of flower abundance values (number of flowers/inflorescences counted in 180 m^2^ per site patch in three rounds) in grasslands from different time-since-fire categories. Relationship between flower abundance values and (b) bare soil cover, %, and (c) vegetation height, cm. FB: freshly-burnt (two to six months after fire; orange circle); IB: intermediate-burnt (about one year after fire; light blue triangle); OB: old-burnt (two years after fire; green square). In a, different letters denote significant differences between categories (p ≤ 0.05) tested with GLMs and Tukey test a posteriori. Boxplots represent the median (solid black line), first and third quartile (box limits), and lines denote maximum and minimum values.

Species composition of plants flowering differed among time-since-fire categories (PERMANOVA F = 1.497, p = 0.01), more specifically between freshly and old-burnt grasslands (p = 0.02). Variation among burned patches from different time-since-fire was mostly related to the first PCoA ordination axis (18%), with freshly-burnt patches associated to positive while old-burnt patches to negative score values (Fig. 7a). The first ordination axis correlated positively with % bare soil (r = 0.80) and temperature (r = 0.42), and negatively with plant biomass (r = -0.54), % moisture (r = -0.57), plant height (r = -0. 64) and % grasses (r = -0.63).

We found seven plant species highly associated (IndVal index > 80%) with grasslands from a specific time-since-fire. Four plant species were associated with freshly, while one species associated with both freshly and intermediate-burnt grasslands. Intermediate and old- burnt grasslands presented one associated plant species each (Table 1).

### Pollinators

We sampled 1,645 insects of 284 species/morphospecies, with 36% bees (67 species), 29% beetles (63 species), 18% flies (70 species), 10% wasps (35 species) and 6% butterflies (49 species) (Fig 1e, f, g, h, i; Table S4 – Supporting Information). Butterfly abundance (χ² = 7.19, df = 2, p = 0.027; Fig. 5a), bee abundance (χ² = 24.32, df = 2, p < 0.001; Fig. 5b), bee richness (χ² = 11.47, df = 2, p = 0.003; Fig. 5c), beetle abundance (χ² = 5.58, df = 2, p = 0.06; Fig. 5d), and beetle richness (df = 2, χ² = 6.89, p = 0.03; Fig. 5e) varied according to time-since-fire. Butterfly abundance was higher in freshly (mean: 24.5 individuals) than in both intermediate (mean: 12.5 individuals) and old-burnt grasslands (mean: 10.7 individuals) (Fig. 5a). Bee abundance and richness were also higher in freshly (mean: 63.5 individuals; mean: 20 species) than in old-burnt (mean: 31.5 individuals, mean: 11 species), but the intermediate patches did not differ from freshly for abundance (mean: 47.7 individuals), and from old for richness (mean: 13 species) (5b and 5c). Coleoptera abundance and richness were higher in old-burnt patches (mean: 55 individuals; mea: 16.2 species) in comparison to intermediate-burnt grasslands (mean: 24 individuals; mean: 9.5 species), but did not differ from freshly-burn (mean: 34.7 individuals; mean: 13.5 species) (Fig. 5d and 5e). Richness of butterflies, and abundance and richness of flies and wasps did not vary with time-since-fire (Table S5 - Supporting Information).

**Fig. 5.**
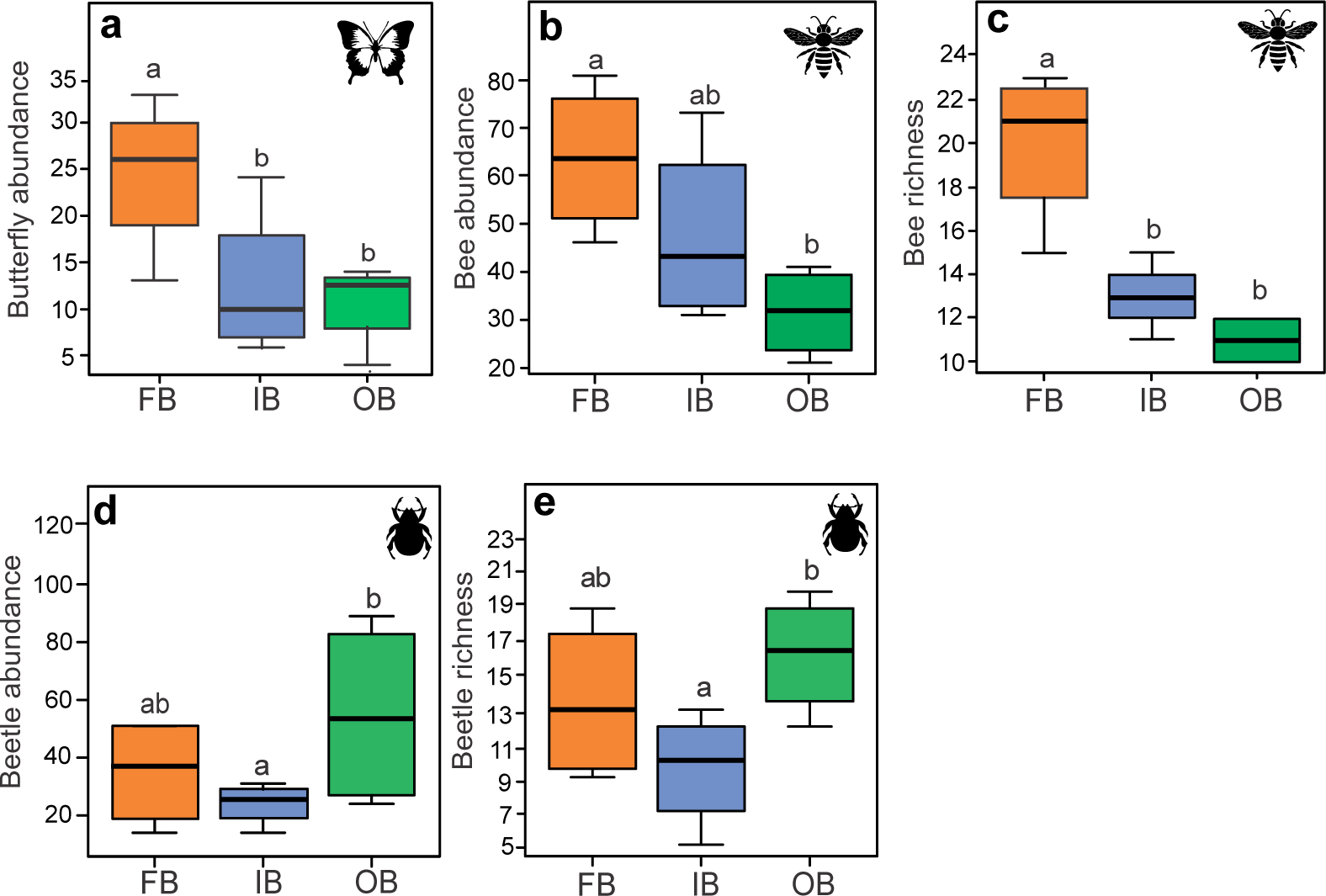
Boxplot of pollinators abundance and richness: (a) butterfly abundance, (b) bee abundance, (c) bee richness, (d) beetle abundance, and (e) beetle richness in grasslands from different time-since-fire categories. FB: freshly-burnt (two to six months after fire; orange); IB: intermediate-burnt (about one year after fire; light blue); OB: old-burnt (two years after fire; green). Different letters denote significant differences between categories (p ≤ 0.05) tested with GLMs and Tukey test a posteriori. Boxplots represent the median (solid black line), first and third quartile (box limits), and lines denote maximum and minimum values. Only significant relationships between pollinators and time-since-fire categories tested with GLMs are shown.

Flower abundance positively explained butterfly abundance (deviance = 8.59, df = 9, p = 0.003, pseudo-R^2^ = 37%; Fig. 6a), bee abundance (deviance = 9.32, df = 9, p= 0.002, pseudo-R^2^ = 41.5%; Fig. 6b), and bee richness (deviance= 11.16, df = 9, p < 0.001, pseudo-R^2^ = 59.8%; Fig. 6c). Vegetation height was a significant predictor of both beetle abundance (deviance = 76.54, df = 8, p < 0.001, pseudo-R^2^= 61.1%; Fig. 6d) and richness (deviance = 76.54, df = 8, p < 0.001, pseudo-R^2^ = 36.2%; Fig. 6e).

**Fig. 6.**
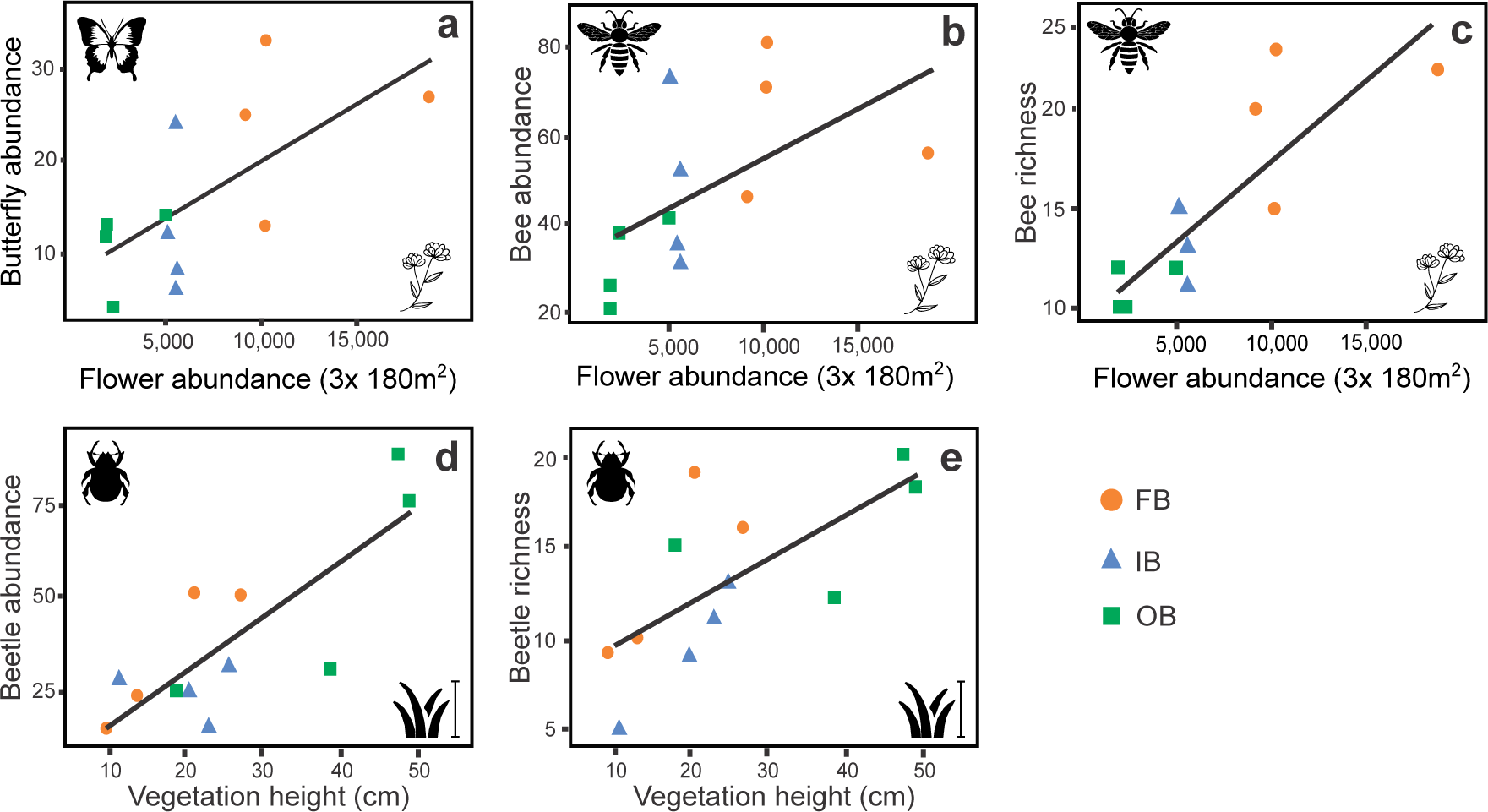
Relationships between flower abundance values (number of flowers/inflorescences counted in 180 m^2^ per patch in three rounds) and (a) butterfly abundance, (b) bee abundance and (c) bee richness. Relationships between vegetation height (cm) and (d) beetle abundance and (g) beetle richness. FB: freshly-burnt (two to six months after fire; orange circle); IB: intermediate- burnt (about one year after fire; light blue triangle); OB: old-burnt (two years after fire; green square). Only significant relationships of pollinators and habitat and flowering plant predictors tested with GLMs are shown.

Bee species composition varied among time-since-fire categories (PERMANOVA F = 1.552, p = 0.02), being significantly different between freshly and old-burnt grasslands (p = 0.05), and intermediate and old-burnt grasslands (p = 0.02). In the PCoA ordination plot, variation among burned patches from different times-since-fire was related to both first (21%) and second (19%) ordination axes (Fig. 7b), with old-burnt patches associated to positive scores in both axes, and freshly and intermediate patches more overdispersed in the ordination. Although the clustering of time-since-fire categories is clear, there was some overlap in the convex hulls enclosing the patches of the given categories. Habitat and plant flowering variables were weakly correlated with the first ordination axes, except plant biomass that presented a strong positive correlation with the first axis (r = 0.70). Plant biomass, plant height, moisture and % grasses were positively related to old-burnt patches, while flower abundance, plant richness, temperature and % bare soil to freshly and intermediate patches (Fig. 7b). Bee and flowering plant composition were significantly correlated (Mantel, r = 0.36; p = 0.01). Species composition of other insect groups did not vary significantly between categories (Table S6 - Supporting Information). We found one bee species highly associated with freshly-burnt grasslands, while one bee and one fly species associated with both freshly and intermediate- burnt grasslands (Table 1).

**Fig. 7.**
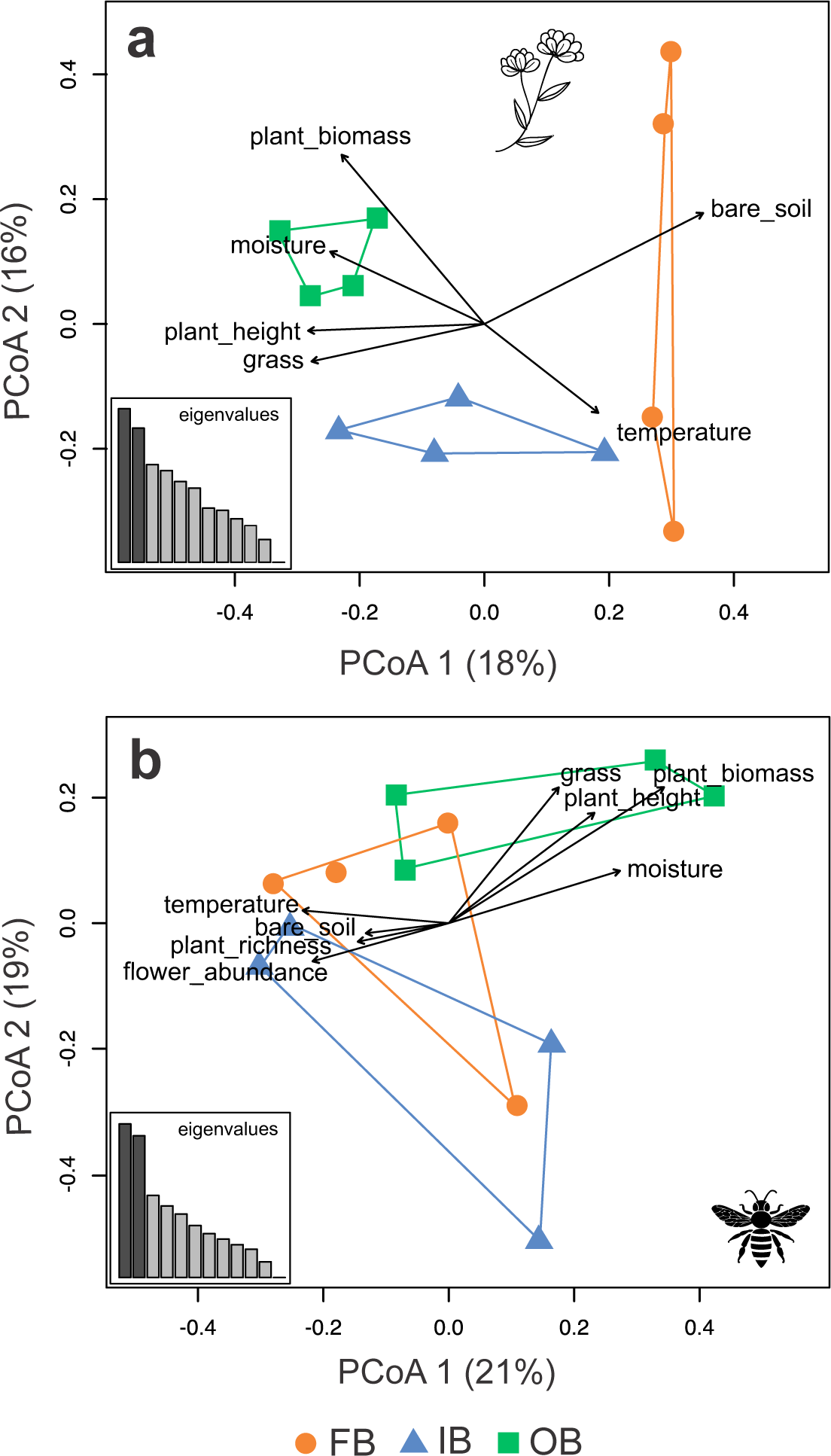
PCoA ordination plots of grassland patches from different time-since-fire categories based on (a) flowering plant and (b) bee community composition. Similarity among patches was measured with Jaccard index for both groups. For each plot, histograms showing the first 12 eigenvalues, and bi-plots of habitat variables (a) and habitat + flowering plant variables (b) are presented. The lengths of the arrows are proportional to their correlation with the axis. FB: freshly-burnt (two to six months after fire; orange circle); IB: intermediate-burnt (about one year after fire; light blue triangle); OB: old-burnt (two years after fire; green square).

## DISCUSSION

In this study, we showed that time-since-fire has a major influence not only on plant communities, but also on pollinators in a grassland ecosystem. Plants increased floral resource concentration in response to fire, and it directly cascaded up to pollinator communities. It is clear that recently burned grasslands benefit an array of flowering plants and pollinators, and that disturbance-suppressed grasslands (e.g., not grazed; not burned for more than one year) present a reduction of floral resources and thereof pollinators decline. Freshly and old-burnt grasslands also have different species compositions. Altogether, our findings have relevant implications for grassland management and biodiversity conservation in non-grazed grasslands of South Brazilian ecosystems.

### Changes in habitat structure

As predicted, grasslands from different times-since-fire greatly differed regarding habitat structure. Freshly-burnt patches have enhanced bare soil, sunlight exposition, and air temperatures, along with decreased moisture, which are driven by the reduction of vegetation aerial biomass (Overbeck *et al*. 2005; Fidelis *et al*. 2012; Podgaiski *et al*. 2014; da Silva *et al*. 2020). Most of this fire-consumed aboveground biomass is in the form of tussock C4 grasses, which are very flammable and represent the dominant plant life-form in grasslands staying longer periods without disturbances (Overbeck *et al*. 2018) (e.g. mean cover in old-burn grasslands was 85%; Fig. 3). The increase in cover of these competitive grasses usually suppresses less competitive species from other plant life forms (e.g., by shading), reducing overall grassland plant diversity (Bond & Keeley 2005; Overbeck *et al*. 2005; Podgaiski *et al*. 2013). Hence, habitat structure resilience to fire in South Brazilian grasslands is high, taking only about two years to affect flowering patterns (Fidelis *et al*. 2012), a time during which tussock grasses, plant biomass and vegetation height notably recover.

### Changes in flower abundance

Associated to this fire-induced habitat dynamics, we found that freshly-burnt grasslands presented a highly increased abundance of flowers, on average four times higher than old-burnt, and twice higher than intermediate-burnt grasslands (Fig. 3), but not an increment in species richness of plants flowering. Flowering stimulated by fire, as showed here, is considered a common trait for herbaceous plants in fire-prone ecosystems worldwide (Byron & Downes 1979; Lamont & Downes 2011; Pyke 2017; Lamont *et al*. 2019). In South Brazil, grassland plant species are known to be well adapted to fire (Overbeck & Pfadenhauer 2007; Fidelis *et al*. 2010a), and Fidelis & Blanco (2014) indeed detected patterns congruent to our findings in nearby grasslands. Most of the plants found flowering in our study site are forbs with large subterranean organs (i.e., bulbs, tuberous roots). These structures accumulate carbohydrates and water, important elements for survival, growth and reproduction (blooming) in environments under constant change from disturbances (Bellingham & Sparrow 2000; Clarke *et al*. 2013; Pausas *et al*. 2018). Thus, shortly after fire, forbs can rapidly resprout and bloom, probably using this stored energy (Fidelis & Blanco 2014; Fidelis *et al*. 2014; Pike 2017). Overall, patterns of flower abundance were directly predicted by the degree of grassland habitat openness (i.e., bare soil exposition), indicating that space availability and reduction of plant competition, especially with dominant tussock grasses, are key drivers encouraging the reproduction of these plants (Fidelis *et al*. 2012; Fidelis & Blanco 2014).

### Changes in pollinator diversity

Little is known about interactions of insect pollinators and plant communities in South Brazilian grasslands, but a few studies have indeed showed butterflies, bees, and beetles as abundant and frequent floral visitors in the study region (Pinheiro *et al*. 2008; Oleques *et al*. 2017; Beal-Neves *et al*. 2020). Bee abundance and richness, and butterfly abundance were greatly amplified in freshly-burnt patches, largely attracted by floral resource improvement in these grasslands. Studies conducted in other regions and ecosystems corroborate these outcomes (Steffan-Dewenter & Tscharntke 2001; Potts et al. 2003b; Collinge *et al*. 2003; Ebeling *et al*. 2008; Dauber *et al*. 2010; Moranz *et al*. 2014, Carbone *et al*. 2019). According to the resource concentration hypothesis (Root 1973; Van Nuland *et al*. 2013), mobile animals are more likely to find and stay longer in resource-dense patches, where they maximize the gain of rewards while minimizing energetic costs of foraging (Charnov 1976). Bees completely rely on floral nectar and pollen for both adult and larval nutrition (Potts *et al*. 2004; Vaudo *et al*. 2015), and constitute one of the pollinator groups most affected by the global pollinator crisis (Bartomeus *et al*. 2019). Adult butterflies depend on nectar for nutrition and breeding, and the fitness of some species can vary according to nectar quality (O’Brien *et al*. 2004; Janz *et al*. 2005; Mevi-Schütz & Erhardt 2005). Besides the attraction to nectar, butterfly abundances may also have increased locally due to the stablishment and growth of their host plants after fire, as showed in other studies (Smallidge & Leopold 1997; Schultz & Crone 1998; Baum & Sharber 2012). However, this was not evaluated directly, deserving detailed exploration in future research. Here we showed that, in the short-term, grassland burnings indirectly aggregated flowers, bees and butterflies. Thus, it could be indicative of the importance of maintenance of these patches in the grassland landscape to ameliorate loss of bee and butterfly pollinators, sustain pollination services, and probably improve flowering plant reproduction (Potts *et al*. 2004; Baum & Sharber 2012; Van Nuland *et al*. 2013).

Flower-visiting beetle communities responded differently than bees and butterflies to time-since-fire. Although freshly and old-burnt grasslands did not differ, vegetation height was a good predictor for beetles, which increased in species and individuals towards taller vegetation of old-burnt grasslands. Vegetation height probably act as a proxy of habitat availability, refuges and oviposition sites for beetles (Dennis *et al*. 1998; Cagnolo *et al*. 2002; Campbell *et al*. 2007). Unlike other pollinators that usually search only for pollen and nectar in flowers, flower-visiting beetle species may be using them as shelter, nest and mating arenas, or even, according to their diverse feeding preferences, as food (e.g., Paulino-Neto 2014). Nevertheless, Coleoptera species are considered primitive pollinators (first pollinators), being of great importance for many angiosperm species reproduction (Bernhardt 2000).

### Changes in plant and bee species composition

Flowering plant community composition varied between freshly and old-burnt grasslands. Among pollinators, bee communities also greatly shifted species between grassland recovery stages, but seemed to follow flowering plants in their composition changes. Fire-induced environmental changes may act as a filtering process, selecting winner and loser species through time (e.g., Pausas & Verdú 2008; Rader *et al*. 2014). The differences in sensitivity to these changes are mostly a result of the traits these species possess (e.g., Pausas & Verdú 2008; Podgaiski *et al*. 2013, 2017). Some species in our study, sharing similar traits, were highly associated to freshly or freshly + intermediate-burnt grasslands. *Galactia marginalis*, *Stevia* sp., *Peltodon longipes*, *Lippia* sp. and *Eryngium pristis*, among the indicator species, all present tuberous roots (Tourn *et al*. 2009; Salimena 2010; Fidelis *et al*. 2014; Gutiérrez *et al*. 2016) which is a trait allowing fast sprout after fire (Fidelis & Blanco 2014; Pausas *et al*. 2018). The bumblebee *Bombus morio* and the sweet bee *Augochlorella acarinata*, also indicators, are generalist pollinators that nest belowground (Michener 2000), and thus need bare ground to excavate their nests, which are very scarce in old-burnt grasslands. Soil condition (i.e., increased bare ground, lower moisture, and warmer temperature), as found recently after fire, has been pointed out as a critical nesting resource for supporting belowground nesting bee communities (Cane & Neff 2011; Buckles & Harmon-Threatt 2019). Moreover, as time since disturbance passes, flowering plant composition shift to more shade-tolerant species or to comparatively better competitors with C4 grasses, e.g., taller herbs, shrubs and sub-shrubs (Ferreira *et al*. 2020), as *Croton gnaphalii*. The establishment of these lignified plants in old-burnt grasslands can potentially provide substrate for a different assemblage of aerial bee nesters (Williams *et al*. 2010). Further studies could complement these results by focusing on trait-based community assembly pattern responses to the fire-induced environmental changes. This approach may help to clarify effects that could not be detected under our species-based models, like for flower-visiting wasp and fly communities (Rader *et al*. 2014).

### Implications and Conclusions

Burning deeply influences grassland species diversity and ecosystem services (Fuhlendorf *et al*. 2009; Podgaiski *et al*. 2013, 2014), and sometimes in very positive ways, beings thus able to improve conservation efforts (Tucker & Cadotte 2013). Here, we contribute to the emergent literature on the fire importance for biodiversity by using a ungrazed grassland in South Brazil as study model. South Brazilian grasslands have already lost more than 50% of their original area especially to agriculture (Cordeiro & Hasenack 2009; Staude *et al*. 2017), and most natural remnants are in private proprieties under extensive grazing. A small percentage of the grassland area is currently under legal protection – which imply disturbance suppression policies (Overbeck *et al*. 2015). The management initiatives on protected areas aim at the suppression of fires and grazing (Pillar & Vélez 2010), which is a strategy based on forest conservation foundations (Bond & Parr 2010; Overbeck *et al*. 2015). Our study shows that total disturbance suppression may lead to flower and pollinator decline (with strong negative impacts on bees), and cause preferential loss of particular species. Consequently, it may not be the best course of action for conservation efforts in such areas. Our work corroborates that a mosaic of grasslands with different times-since-fire (i.e., high in pyrodiversity; Ponisio *et al*. 2016; Parr & Brockett 1999) associated to adjacent forest physiognomies should be highly valued for conservation purposes (Luza *et al*. 2014).

Lastly, it is worth mentioning again that fire effects on ecosystems and their biodiversity are usually complex, and greatly tied to specific fire regime characteristics, including ecosystem fire history, fire frequency, intensity and burned area size (Driscoll *et al*. 2010; Bowman *et al*. 2011). Further understanding on how these variables associated with burning affect flowering plants, pollinators, and interaction networks is essential for guiding future management actions. Here, we advanced on time-since-fire effects only, pointing out the role of freshly-burnt patches as high quality, resource-rich habitats for pollinators, indirectly contributing to pollination services (Ponisio *et al*. 2016; Brown *et al*. 2017; Peralta *et al*. 2017).

