## Supplemental Material for "Burning for grassland pollination: recently burned patches promote plant flowering and insect pollinators"

**Table S1.** Description of the 12 studied grassland patches at Parque Natural Municipal Saint’ Hilaire. Viamão municipality. Rio Grande do Sul State. Brazil. Sampling season A: 2015. November–2016. April; B: 2016. November–2017. April. Patch size represents the approximate size of the grassland burned area. Distance to urban area represents the linear distance from the center of the patch to the border of the nearest urban area. Percentage of forest and grasslands available in a buffer of 300 m radius around the center of the patch. sd = standard deviation. Freshly-burnt: two to six months; Intermediate-burnt: about one year; Old-burnt: two years after fire.

| Grassland site | Sampling season | Time-since-fire category | Size (ha) | Distance to urban area (m) | % grassland | % forest |
| --- | --- | --- | --- | --- | --- | --- |
| 1 | A | Freshly | 0.80 | 129 | 0.25 | 0.66 |
| 2 | A | freshly | 0.60 | 1,082.36 | 0.41 | 0.59 |
| 3 | A | intermediate | 1.27 | 320.58 | 0.53 | 0.46 |
| 4 | A | intermediate | 7.74 | 757.63 | 0.49 | 0.50 |
| 5 | A | old | 2.55 | 626.96 | 0.25 | 0.74 |
| 6 | A | old | 1.10 | 167.81 | 0.23 | 0.67 |
| 7 | B | freshly | 3.86 | 547.26 | 0.44 | 0.56 |
| 8 | B | freshly | 2.92 | 255.18 | 0.60 | 0.32 |
| 9 | B | intermediate | 1.71 | 227.51 | 0.49 | 0.48 |
| 10 | B | intermediate | 2.33 | 888.15 | 0.37 | 0.52 |
| 11 | B | old | 1.64 | 144.38 | 0.21 | 0.65 |
| 12 | B | old | 4.49 | 879.74 | 0.46 | 0.54 |
| Average (sd) | | freshly | 2.04 (1.60) | 503.45 (0.04) | 0.42 (0.14) | 0.53 (0.15) |
| Average (sd) | | intermediate | 3.26 (3.02) | 548.46 (0.01) | 0.47 (0.07) | 0.49 (0.02) |
| Average (sd) | | old | 2.44 (1.49) | 454.72 (0.06) | 0.29 (0.11) | 0.65 (0.08) |

**Table S2-** VIF values for predictor variables in the final models.

| **Full model predictors** | **Reduced models** | **final VIF** |
| --- | --- | --- |
| Plant descriptor ~ biomass + plant height +bare soil + grass | Flower abundance ~ plant height + bare soil | 1.55 |
| Pollinator descriptor ~ flower abundance+biomass +plant height +bare soil + grass | Beetle abundance ~ flower abundance + plant height | 1.47 |
|  | Beetle richness ~ flower abundance + plant height | 1.50 |
|  | Bee abundance ~ flower abundance + plant height | 1.30 |
|  | Bee richness ~ flower abundance + plant height | 1.20 |
|  | Butterfly abundance ~ flower abundance + plant height | 1.20 |
|  | Butterfly richness ~ flower abundance + plant height | 1.27 |

**Table S3.** Species composition of plants in grasslands from different time-since-fire categories. Freshly-burnt: two to six months; Intermediate-burnt: about one year; Old-burnt: two years after fire.

| **Family** | **Plant species** | **Freshly-burnt** | **Intermediate-burnt** | **Old-burnt** |
| --- | --- | --- | --- | --- |
| Acanthaceae | *Justicia axillaris* (Nees) Lindau | x |  |  |
| Amaranthaceae | *Pfaffia tuberosa* (Spreng.) Hicken | x | x | x |
| Apiaceae | *Eryngium ciliatum* Cham. & Schltdl. | x |  |  |
|  | *Eryngium elegans* Cham. Et Schlecht. | x | x | x |
|  | *Eryngium eriophorum* Cham. & Schltdl. | x | x |  |
|  | *Eryngium horridum* Malme | x | x | x |
|  | *Eryngium pristis* Cham. & Schltdl. | x | x |  |
|  | *Eryngium sanguisorba* Cham. Et Schlecht. | x | x | x |
| Apocynaceae | *Asclepias mellodora* A. St.-Hil. | x |  |  |
|  | *Mandevilla coccinea* (Hook. & Arn.) Woodson |  | x |  |
|  | *Mandevilla longiflora* (Desf.) Pichon | x |  |  |
|  | *Oxypetalum arnottianum* H. Buek | x |  |  |
| Aristolochiaceae | *Aristolochia sessilifolia (*Klotzsch) Duch. | x |  | x |
| Asteraceae | *Achyrocline satureioides* (Lam.) DC. |  |  | x |
|  | *Acmella bellidioides* (Smith in Rees) R.K. Jansen | x |  |  |
|  | *Aspilia montevidensis* (Spreng.) Kuntze | x | x | x |
|  | *Baccharis articulata* (Lam.) Pers. |  | x |  |
|  | *Baccharis leucopappa* DC. |  |  | x |
|  | *Baccharis megapotamica* Spreng | x |  |  |
|  | *Baccharis pentodonta* Malme |  | x |  |
|  | *Baccharis spicata* (Lam.) Baill. |  |  | x |
|  | *Baccharis trimera* (Less.) DC. |  |  | x |
|  | *Calea uniflora* Less. | x |  | x |
|  | *Campuloclinium macrocephalum* (Less.) DC. | x |  | x |
|  | *Chaptalia runcinata* Kunth | x |  |  |
|  | *Chromolaena ascendens* (Sch.Bip. ex Baker) R.M.King & H.Rob. | x | x | x |
|  | *Chromolaena congesta* (Hook. & Arn.) R.M.King & H.Rob. | x | x | x |
|  | *Chromolaena hirsuta* (Hook. & Arn.) R.M.King & H.Rob. | x | x | x |
|  | *Chromolaena squarrulosa* (Hook. & Arn.) R.M.King & H.Rob. | x | x | x |
|  | *Chrysolaena flexuosa* (Sims) H. Rob. | x | x | x |
|  | *Conyza lorentzii* Griseb. | x |  | x |
|  | *Conyza primulifolia* (Lam.) Cuatrec. & Lourteig | x | x |  |
|  | *Criscia stricta* (Spreng.) L. Katinas |  |  | x |
|  | *Disynaphia ligulifolia* (Hook. & Arn.) R.M.King & H.Rob. | x | x | x |
|  | *Eupatorium tanacetifolium* Gill. ex Hook. & Arn. | x | x | x |
|  | *Gamochaeta* sp. |  |  | x |
|  | *Lessingianthus hypochaeris* (DC.) H.Rob. | x | x |  |
|  | *Moquiniastrum cordatum* (Less.) G. Sancho |  | x |  |
|  | *Orthopappus angustifolius* (Sw.) Gleason | x | x | x |
|  | *Porophyllum curticeps* Malme |  |  | x |
|  | *Porophyllum lanceolatum* DC. | x |  | x |
|  | *Pterocaulon rugosum (*Vahl) Malme | x | x | x |
|  | *Stenachaenium megapotamicum* (Spreng.) Baker | x | x |  |
|  | *Stenocephalum megapotamicum* (Spreng.) Sch.Bip. | x | x | x |
|  | *Stevia* sp. | x | x |  |
|  | *Vernonia flexuosa* Sims |  | x | x |
|  | *Vernonia nudiflora* Less. | x | x | x |
|  | *Vernonia squarrosa* Less. (Less) | x |  |  |
| Bromeliaceae | *Dyckia leptostachya* Baker |  | x | x |
| Campanulaceae | *Wahlenbergia linarioides* (Lam.) A.DC. |  |  | x |
| Convolvulaceae | *Cuscuta xanthochortos* Mart. |  |  | x |
|  | *Evolvulus sericeus* Sw. | x | x | x |
| Euphorbiaceae | *Croton gnaphalii* Baill. |  | x | x |
|  | *Euphorbia selloi* (Klotzsch & Garcke) Boiss. | x | x | x |
| Fabaceae | *Aeschynomene falcata* (Poir.) DC. | x | x | x |
|  | *Centrosema virginianum* (L.) Benth. |  | x |  |
|  | *Chamaecrista nictitans* (L.) Moench | x | x | x |
|  | *Chamaecrista repens* (Vogel) H.S. Irwin & Barneby | x | x | x |
|  | *Collaea stenophylla* (Hook et Arn.) Benth. |  |  | x |
|  | *Crotalaria tweediana* Benth. | x |  | x |
|  | *Desmanthus tatuhyensis* Hoehne | x | x | x |
|  | *Desmodium barbatum* (L.) Benth. |  |  | x |
|  | *Galactia marginalis* Benth. ex Benth. & Hook. f. | x |  |  |
|  | *Galactia pretiosa* Burkart | x |  |  |
|  | *Macroptilium prostratum* (Benth.) Urb. | x | x | x |
|  | *Rhynchosia corylifolia* Mart. ex Benth. | x |  |  |
|  | *Rhynchosia diversifolia* Micheli | x | x | x |
|  | *Stylosanthes leiocarpa* Vogel |  | x | x |
|  | *Stylosanthes montevidensis* Vogel | x | x | x |
|  | *Zornia* sp. | x | x | x |
| Iridaceae | *Buchnera longifolia* Kunth |  |  | x |
|  | *Cypella amplimaculata* Chauveau & L.Eggers. A–C. | x |  | x |
|  | *Cypella pusilla* (Link & Otto) Benth. & Hook. f. |  | x |  |
|  | *Herbertia pulchella* Sweet | x |  |  |
|  | *Sisyrinchium sellowianum* Klatt | x |  | x |
|  | *Sisyrinchium vaginatum* Spreng. | x | x | x |
| Lamiaceae | *Glechon ciliata* Benth. | x | x | x |
|  | *Peltodon longipes* A.St.-Hil. ex Benth. | x | x | x |
| Lythraceae | *Cuphea glutinosa* Cham. & Schltdl. | x | x | x |
| Malpighiaceae | *Galphimia australis* Chodat |  | x |  |
| Malvaceae | *Pavonia hastata* Cav. |  |  | x |
|  | *Sida* sp. | x | x | x |
|  | *Wissadula glechomifolia* (A. St.-Hil.) R.E. Fr. | x | x |  |
| Melastomataceae | *Tibouchina gracilis* (Bonpl.) Cogn. | x | x | x |
| Myrtaceae | *Campomanesia aurea* O.Berg | x | x | x |
|  | *Eugenia dimorpha* O.Berg |  |  | x |
| Orchidaceae | *Sacoila lanceolata* (Aubl.) Garay |  | x |  |
| Orobanchaceae | *Agalinis communis* (Cham. & Schltdl.) D'Arcy |  | x | x |
|  | *Buchnera longifolia* Kunth | x | x | x |
|  | *Castilleja arvensis* Schltdl. & Cham | x |  |  |
| Passifloraceae | *Passiflora foetida* L. |  |  | x |
|  | *Piriqueta taubatensis* (Urb.) Arbo | x |  |  |
| Plantaginaceae | *Angelonia integerrima* Sprengel | x |  |  |
| Polygalaceae | *Polygala adenophylla* A. St.-Hill. & Moq. | x | x |  |
|  | *Polygala* sp. | x | x | x |
| Rubiaceae | *Borreria capitata* (Ruiz & Pav.) DC. | x | x | x |
|  | *Diodella apiculata* (Willd. ex Roem. & Schult.) Delprete |  |  | x |
|  | *Galianthe fastigiata* Griseb. | x | x | x |
|  | *Richardia grandiflora* (Cham. & Schltdl.) Steud. | x | x | x |
|  | *Spermacoce verticillata* L. | x | x | x |
| Solanaceae | *Calibrachoa ovalifolia* (Miers) Stehmann & Semir | x | x |  |
| Verbenaceae | *Glandularia marrubioides* (Cham.) Tronc. | x | x |  |
|  | *Glandularia thymoides* (Cham.) N. O' Leary |  |  | x |
|  | *Lippia* sp. | x | x |  |
|  | *Verbena litoralis* Kunth. | x | x | x |

**Table S4.** Species composition of insects visiting flowers in grasslands from different time-since-fire categories. Freshly-burnt: two to six months; Intermediate-burnt: about one year; Old-burnt: two years after fire.

| **Order** | **Flower visitor species** | **Freshly -burnt** | **Intermediate- burnt** | **Old-burnt** |
| --- | --- | --- | --- | --- |
| Coleoptera | *Agrilus sacer tremolerasi* (Obenberger. 1933) | x | x | x |
|  | *Agrilus* sp. 1 |  |  | x |
|  | Alticini sp.1 |  |  | x |
|  | Apioninae sp. 1 | x | x | x |
|  | Apioninae sp. 2 | x | x | x |
|  | Apioninae sp. 3 |  |  | x |
|  | *Astylus* sp. 1 |  |  | x |
|  | *Astylus* sp. 2 |  |  | x |
|  | *Atelopteryx compsoceroides* (Lacordaire. 1869) |  |  | x |
|  | Baridinae sp. 1 |  | x | x |
|  | Baridinae sp. 2 | x | x | x |
|  | Baridinae sp. 2A | x | x | x |
|  | Baridinae sp. 3 | x | x | x |
|  | Baridinae sp. 3A | x |  | x |
|  | Baridinae sp. 4 | x | x | x |
|  | Baridinae sp. 4A | x |  |  |
|  | Baridinae sp. 5 | x |  | x |
|  | *Brasilaphthona hortensia* (Bechyné 1955) | x | x | x |
|  | Bruchinae sp. 1 | x |  | x |
|  | Bruchinae sp.2 | x |  |  |
|  | Bruchinae sp. 3 | x |  | x |
|  | Buprestidae sp. 1 |  |  | x |
|  | Buprestidae sp. 2 |  |  | x |
|  | Cantharidae sp. 1 | x | x | x |
|  | Cantharidae sp. 2 | x |  | x |
|  | Cantharidae sp. 3 |  |  | x |
|  | *Chalcodermus humeridens* (Faust. 1894) |  | x |  |
|  | *Chalcodermus* sp. 1 | x |  |  |
|  | *Chauliognathus scriptus* (Germar. 1824) | x | x |  |
|  | *Chauliognathus* sp. 1 | x | x | x |
|  | Clytrini sp. 1 | x |  | x |
|  | Clytrini sp. 2 | x |  | x |
|  | Clytrini sp. 3 |  |  | x |
|  | *Cosmisoma brullei* (Mulsant. 1863) | x | x | x |
|  | Cryptocephalinae sp. 1 |  | x |  |
|  | *Cryptocephalus* sp. 1 | x |  |  |
|  | Curculionidae sp. 1 |  |  | x |
|  | *Diabrotica* sp. 1 | x |  |  |
|  | *Diabrotica* sp. 2 |  |  | x |
|  | *Diabrotica* sp. 3 |  | x |  |
|  | *Discodon* sp. 1 | x | x | x |
|  | *Disonycha* sp.1 | x |  |  |
|  | *Enoclerus* sp. 1 | x |  |  |
|  | Eumolpinae sp. 1 |  | x | x |
|  | *Lasionota bonaerensis* (Moore. 1997) |  |  | x |
|  | *Lasionota fairmairei* (Kerremans. 1897) |  |  | x |
|  | *Lasionota* sp. 1 | x | x |  |
|  | *Lexiphanes biplagiatus* (Boheman. 1848) | x |  | x |
|  | *Lexiphanes* sp. 1 |  |  | x |
|  | *Lobaspis squamosus* (Boheman. 1836). |  |  | x |
|  | *Macraspis dichroa* (Mannerheim. 1829) |  |  | x |
|  | Molytinae sp. 1 |  |  | x |
|  | Molytinae sp. 2 |  |  | x |
|  | Mordellidae sp. 1 | x |  | x |
|  | Mordellidae sp. 2 | x | x | x |
|  | Naupactini sp. 1 | x |  |  |
|  | Naupactini sp. 2 |  |  | x |
|  | Naupactini sp. 3 |  |  | x |
|  | *Odontocera flavicauda* (Bates. 1873) | x |  |  |
|  | *Paranapiacaba* sp. 1 |  |  | x |
|  | Phalacridae sp.1 |  |  | x |
|  | *Pristimerus* sp. 1 |  |  | x |
|  | *Spintherophyta* sp. 1 | x | x | x |
| Diptera | *Anthomyia* sp. 1 | x |  |  |
|  | Asilidae sp. 1 |  | x |  |
|  | Asilidae sp. 2 |  |  | x |
|  | Bibionidae sp. 1 |  | x |  |
|  | Bibionidae sp. 2 |  | x |  |
|  | Bombyliidae sp.1 | x |  | x |
|  | Bombyliidae sp. 2 |  | x |  |
|  | Bombyliidae sp. 3 | x |  |  |
|  | Bombyliidae sp. 4 |  | x |  |
|  | Bombyliidae sp. 5 |  | x |  |
|  | Bombyliidae sp. 6 |  |  | x |
|  | Calliphoridae sp. 1 |  |  | x |
|  | *Chryrsomia albiceps* (Wiedemann. 1819) |  |  | x |
|  | Curtonotidae sp. 1 | x | x | x |
|  | Diptera sp. 1 |  | x |  |
|  | Empididae sp. 1 |  | x |  |
|  | Ephydridae sp. 1 | x |  |  |
|  | *Hemilucilia semidiaphana* (Rondani. 1850) |  | x |  |
|  | Lonchaeidae sp. 1 |  |  | x |
|  | *Lucilia eximia* (Wiedemann. 1819) |  | x |  |
|  | *Lucilia* sp. 1 | x |  |  |
|  | *Musca domestica* (Linnaeus. 1758) |  |  | x |
|  | *Ornidia obesa* (Fabricius. 1775) | x |  |  |
|  | *Pseudodorus clavatus* (Fabricius. 1794) | x | x | x |
|  | Sarcophagidae sp. 1 | x | x |  |
|  | Sarcophagidae sp. 2 | x | x | x |
|  | Sarcophagidae sp. 3 | x |  | x |
|  | Sarcophagidae sp. 4 | x | x |  |
|  | Sarcophagidae sp. 5 | x | x |  |
|  | Sarcophagidae sp. 6 |  |  | x |
|  | Sarcophagidae sp. 7 |  |  | x |
|  | Sarcophagidae sp. 8 | x |  | x |
|  | Sarcophagidae sp. 9 |  |  | x |
|  | Sarcophagidae sp. 10 | x |  |  |
|  | Sarcophagidae sp. 11 |  | x |  |
|  | Sarcophagidae sp. 12 | x |  |  |
|  | Sarcophagidae sp. 13 |  | x |  |
|  | Sarcophagidae sp. 14 | x | x |  |
|  | Sarcophagidae sp. 15 |  |  | x |
|  | Sarcophagidae sp. 16 |  | x |  |
|  | Stratiomyidae sp. 1 |  | x |  |
|  | Stratiomyidae sp. 2 |  | x |  |
|  | *Systropus* sp. 1 |  |  | x |
|  | Syrphidae sp. 1 |  |  | x |
|  | Syrphidae sp. 2 | x |  | x |
|  | Syrphidae sp. 3 |  |  | x |
|  | Syrphidae sp. 4 | x | x | x |
|  | Syrphidae sp. 5 | x | x | x |
|  | Syrphidae sp. 6 | x | x |  |
|  | Syrphidae sp. 7 | x | x | x |
|  | Syrphidae sp. 8 | x | x | x |
|  | Syrphidae sp. 9 |  | x |  |
|  | Tabanidae sp. 1 |  | x | x |
|  | Tabanidae sp. 2 |  | x |  |
|  | Tabanidae sp. 3 | x | x | x |
|  | Tachinidae sp. 1 | x |  |  |
|  | Tachinidae sp. 2 |  | x |  |
|  | Tachinidae sp. 3 |  | x |  |
|  | Tachinidae sp. 4 |  | x |  |
|  | Tachinidae sp. 5 |  |  | x |
|  | Tachinidae sp. 6 | x | x | x |
|  | Tachinidae sp. 7 |  |  | x |
|  | Tachinidae sp. 8 |  | x |  |
|  | Tachinidae sp. 9 | x | x |  |
|  | Tachinidae sp. 10 |  | x |  |
|  | Tachinidae sp. 11 |  |  | x |
|  | *Tricopoda* sp. 1 | x |  |  |
|  | *Trupanea* sp.1 | x |  |  |
|  | *Ulidiida*e sp. 1 |  | x |  |
|  | *Xanthaciura* sp. 1 |  |  | x |
| Hymenoptera (bees) | Anthidiini sp. 1 | x |  |  |
|  | Anthrenoides sp. 1 | x |  |  |
|  | *Apis mellifera* (Linnaeus. 1758) | x | x | x |
|  | *Augochlora (Augochlora) amphitrite* (Schrottky. 1909) | x | x |  |
|  | *Augochlora (Oxystoglossella) iphigenia* (Holmberg. 1886) | x | x |  |
|  | *Augochlora cfr. foxiana* (Cockerell. 1900) | x |  |  |
|  | *Augochlora* sp. 1 |  |  | x |
|  | *Augochlora* sp. 2 |  |  | x |
|  | *Augochlorella acarinata* (Coelho. 2004) | x | x |  |
|  | *Augochlorella ephyra* (Schrottky. 1910) |  | x | x |
|  | *Augochloropsis anisitsi* (Schrottky. 1908) |  | x |  |
|  | *Augochloropsis multiplex* (Vachal. 1903) |  | x | x |
|  | *Augochloropsis* sp. 1 | x | x | x |
|  | *Augochloropsis* sp. 2 | x |  |  |
|  | *Augochloropsi*s sp. 3 | x |  |  |
|  | *Augochloropsis* sp. 4 | x | x | x |
|  | *Bombus (Fervidobombus) morio* (Swederus. 1787) | x |  |  |
|  | *Centris* sp. 1 |  | x |  |
|  | *Centris* sp. 2 | x | x |  |
|  | *Centris* sp. 3 | x |  |  |
|  | *Centris* sp. 4 | x |  |  |
|  | *Ceratina* (*Crewella*) sp. 1 | x | x | x |
|  | *Ceratina* (*Crewella*) sp. 2 | x | x |  |
|  | *Ceratina (Neoclavicera) asunciana* (Strand. 1910) | x | x | x |
|  | *Ceratina (Rhysoceratina) volitans* (Schrottky. 1907) |  |  | x |
|  | *Dialictus* sp. 1 | x | x |  |
|  | *Dialictus* sp. 2 |  |  | x |
|  | *Dialictus* sp. 3 | x | x | x |
|  | *Eucerini* sp. 1 | x |  |  |
|  | *Eucerini* sp. 2 | x |  |  |
|  | *Eucerini* sp. 3 | x |  |  |
|  | *Eufriesea* sp. 1 |  | x |  |
|  | *Exomalopsis* sp. 1 | x |  | x |
|  | *Exomalopsis* sp. 2 | x | x |  |
|  | *Gaesischia* sp. 1 | x |  |  |
|  | *Gaesischia* sp. 2 | x | x |  |
|  | *Gaesischia* sp. 3 | x |  |  |
|  | *Gaesischia* sp. 4 |  | x |  |
|  | *Gaesischia* sp. 5 | x |  |  |
|  | *Megachile* sp. 1 | x |  | x |
|  | *Megachile* sp. 2 | x | x |  |
|  | *Megachile* sp. 3 |  | x |  |
|  | *Megachile* sp. 4 |  |  | x |
|  | *Megachil*e sp. 5 | x | x | x |
|  | *Megachile* sp. 6 | x |  |  |
|  | *Megachile* sp. 7 |  | x |  |
|  | *Megachile* sp. 8 | x |  |  |
|  | *Megachile* sp. 9 |  | x |  |
|  | *Melissoptila* sp. 1 | x |  | x |
|  | *Plebeia* sp. 1 | x | x | x |
|  | *Plebeia* sp. 2 | x |  |  |
|  | *Psaenythia* sp. 1 |  |  | x |
|  | *Psaenythia* sp. 2 | x | x | x |
|  | *Psaenythia* sp. 3 | x |  |  |
|  | *Pseudaugochlora graminea* (Fabricius. 1804) |  |  | x |
|  | *Ptilothrix* sp. 1 |  |  | x |
|  | *Scaptotrigona bipunctata* (Lepeletier. 1836) | x |  |  |
|  | *Schwarziana* sp. 1 | x | x | x |
|  | *Tetragonisca angustula* (Latreille. 1811) | x | x | x |
|  | *Thectochlora hamata* (Gonçalves & Melo. 2006) | x |  |  |
|  | *Thygater* sp .1 | x |  |  |
|  | *Trigona spinipes* (Fabricius. 1793) | x | x |  |
|  | *Xylocopa (Neoxylocopa) augusti* (Lepeletier. 1841) | x |  | x |
|  | *Xylocopa (Xylocospila) bambusae* (Schrottky. 1902) |  |  | x |
|  | *Xylocopa (Nanoxylocopa) ciliata* (Burmeister. 1876) | x | x | x |
|  | *Xylocopa (Neoxylocopa) frontalis* (Olivier. 1789) | x | x |  |
|  | *Xylocopa (Neoxylocopa) nigrocincta* (Smith. 1854) | x |  |  |
| Hymenoptera (wasps) | *Bembix* sp. 1 |  | x |  |
|  | *Brachygastra lecheguana* (Latreille. 1824) |  | x | x |
|  | Cabronidae sp. 1 | x | x | x |
|  | Cabronidae sp. 2 | x |  |  |
|  | Cabronidae sp. 3 |  | x |  |
|  | Cabronidae sp. 4 |  |  | x |
|  | Cabronidae sp. 5 |  |  | x |
|  | *Cyphomenes* sp. 1 |  | x |  |
|  | *Hemipepsis* sp. 1 |  | x | x |
|  | *Hypalastoroides paraguayensis* (Zavattari. 1911) | x |  |  |
|  | *Minixi* sp. 1 | x |  | x |
|  | *Minixi* sp. 2 |  |  | x |
|  | *Mischocyttarus drewseni* (Saussure. 1857) | x | x | x |
|  | *Montzumia infernalis* (Spinola. 1851) | x |  |  |
|  | *Omicron aurantiopictum* (Soika. 1978) | x |  | x |
|  | *Pachodynerus* sp. 1 |  |  | x |
|  | *Pachodynerus* sp. 2 |  |  | x |
|  | P*achodynerus* sp. 3 |  | x |  |
|  | *Pachymenes* sp. 1 | x |  |  |
|  | *Pachymenes* sp. 2 |  |  | x |
|  | *Pachyminixi* sp. 1 |  | x |  |
|  | *Pepsis* sp. 1 |  | x |  |
|  | *Pepsis* sp. 2 |  | x |  |
|  | *Pepsis* sp. 3 | x | x |  |
|  | *Pirhosigma cf. deforme* (Fox. 1899) |  |  | x |
|  | *Pirhosigma* sp. 1 |  | x |  |
|  | *Polistes billardieri* (Fabricius. 1804) | x | x | x |
|  | *Polistes carnifex* (Fabricius. 1775) |  |  | x |
|  | *Polybia ignobilis* (Haliday. 1836) | x | x | x |
|  | *Polybia sericea* (Olivier. 1791) |  | x | x |
|  | *Polybia scutellaris* (White. 1841) |  | x | x |
|  | *Sceliphron* sp. 1 |  | x |  |
|  | Sphecidae sp. 1 | x |  |  |
|  | Sphecidae sp. 2 | x |  |  |
|  | *Zeta argillaceum* (Linnaeus. 1758) | x |  | x |
| Lepidoptera | *Achlyodes mithridates* (Fabricius. 1793) |  | x |  |
|  | *Actinote* sp. | x |  | x |
|  | *Agraulis vanillae maculosa* (Linnaeus. 1758) | x | x | x |
|  | *Apodemia castanea* (Prittwitz. 1865) | x |  |  |
|  | *Aricoris gauchoana* (Stichel. 1910) |  | x | x |
|  | *Aricoris monotona* (Stichel. 1910) | x |  |  |
|  | *Aricoris notialis* (Stichel. 1910) | x |  |  |
|  | *Battus polydamas* (Linnaeus. 1758) | x |  |  |
|  | *Chioides catillus* (Cramer. 1779) | x | x | x |
|  | *Cogia cf. hassan evansi* (Bell. 1937) | x | x |  |
|  | *Contrafacia muattina* (Schaus. 1902) |  |  | x |
|  | *Danaus plexippus* (Linnaeus. 1758) |  |  | x |
|  | *Dryas iulia* (Fabricius. 1775) | x | x | x |
|  | *Emesis russula* (Stichel. 1910) |  | x |  |
|  | *Euptoieta hortensia* (Blanchard. 1852) | x |  | x |
|  | *Euryades corethrus* (Boisduval. 1836) | x |  |  |
|  | *Eurema deva* (Cramer. 1780) |  |  | x |
|  | *Eurema elathia* (Cramer. 1777) | x | x |  |
|  | *Heliopetes arsalte* (Linnaeus. 1758) | x |  | x |
|  | *Hemiargus hanno* (Stoll. 1790) | x | x | x |
|  | *Heraclides anchisiades* (Esper. 1788) |  | x |  |
|  | *Heraclides hectorides* (Esper. 1794) | x |  |  |
|  | Hesperiidae sp. 1 | x | x | x |
|  | Hesperiidae sp. 2 | x | x |  |
|  | Hesperidae sp. 3 |  |  | x |
|  | Hesperidae sp. 4 |  |  | x |
|  | *Hypanartia bella (*Fabricius. 1793) |  | x |  |
|  | *Junonia evarete* (Cramer. 1779) | x | x | x |
|  | *Lerodea eufala eufala* (Edwards. 1869) |  |  | x |
|  | *Mechanitis l. lysimnia* (Fabricius. 1793) | x |  |  |
|  | *Nicolaea torris* (Druce. 1907) | x |  |  |
|  | Nymphalidae sp. 1 | x |  |  |
|  | *Ortilia dicoma* (Hewitson. 1864) |  | x |  |
|  | *Pampasatyrus periphas* (Godart. 1824) |  |  | x |
|  | *Panoquina hecebolus* (Scudder. 1872) | x |  |  |
|  | *Phoebis sennae* (Linnaeus. 1758) |  | x | x |
|  | *Pseudolucia parana* (Bálint. 1993) |  |  | x |
|  | *Pyrgus orcus* (Stoll. 1780) | x |  | x |
|  | *Rekoa palegon* (Cramer. 1780) |  | x |  |
|  | *Strymon bazochii* (Godart. 1824) |  |  | x |
|  | *Strymon rana* (Schaus. 1902) | x |  |  |
|  | *Tegosa orobia* (Hewitson. 1864) | x | x |  |
|  | *Urbanus dorantes* (Stoll. 1790) |  | x | x |
|  | *Urbanus evenus* (Ménétriés. 1855) | x | x | x |
|  | *Urbanus simplicius* (Stoll. 1790) |  | x |  |
|  | *Urbanus teleus* (Hübner. 1821) | x | x | x |
|  | *Vanessa braziliensis* (Moore. 1883) | x | x | x |
|  | *Vanessa myrinna* (Doubleday. 1849) | x |  |  |
|  | *Wallengrenia premnas* (Wallengren. 1860) | x | x | x |

**Table S5.** Likelihood-ratio test results comparing negative-binomial models with time-since-fire categories as predictor (dependent variable ~ season+ time-since-fire categories) with null models (dependent variable ~ season) for flowering plant richness and pollinator descriptors. Only models that did not differ (p>0.05) are shown.

| **Dependent variable** | **df** | **Chi squared** χ² | **p-value** |
| --- | --- | --- | --- |
| Flowering plant richness | 2 | 3.59 | 0.16 |
| Butterfly richness | 2 | 4.45 | 0.10 |
| Fly abundance | 2 | 1.67 | 0.43 |
| Fly richness | 2 | 0.99 | 0.61 |
| Wasp abundance | 2 | 2.39 | 0.30 |
| Wasp richness | 2 | 0.37 | 0.83 |

**Table S6.** PERMANOVA test results (9999 permutations) showing non-significant (p>0.05) relationships between insect pollinator composition and time-since-fire categories. Jaccard similarity index was used.

| **Dependent variable** | **PERMANOVA** |
| --- | --- |
| Butterfly composition | F= 1.01; p = 0.44 |
| Fly composition | F=0.91; p = 0.60 |
| Beetle composition | F= 0.85; p= 0.68 |
| Wasp composition | F= 0.93; p = 0.58 |
